# Cross-species blastocyst chimerism between nonhuman primates using iPSCs

**DOI:** 10.1101/635250

**Authors:** Morteza Roodgar, Fabian P. Suchy, Vivek Bajpai, Jose G. Viches-Moure, Joydeep Bhadury, Angelos Oikonomopoulos, Joseph C. Wu, Joseph L. Mankowski, Kyle M. Loh, Hiromitsu Nakauchi, Catherine A. VandeVoort, Michael P. Snyder

## Abstract

Through the production of chimeric animals, induced pluripotent stem cells (iPSCs) can generate personalized organs with diverse applications for both basic research and translational medicine. This concept was first validated in rodents by forming a rat pancreas in mice and vice versa. However, the potential use of human iPSCs to generate xenogenic organs in other species is technically and ethically difficult. Recognizing these concerns, we explored the generation of chimeric nonhuman primates (NHP) embryos, by injecting either chimpanzee or pig-tailed macaque iPSCs into rhesus macaque embryos. We first derived iPSCs from chimpanzees and pig-tailed macaques. We found that the chimpanzee iPSCs mixed well with human iPSCs during *in vitro* co-culture and differentiation. The differentiation of mixed human and chimpanzee iPSCs formed functioning cardiomyocyte layers in vitro, whereas human or chimpanzee iPSC mixed with pig-tailed macaque or mouse cells do not; these results indicate that chimpanzee and human cells are closely related in function. Considering the ethical aspects of injecting human iPSCs into nonhuman primate blastocysts, we tested whether chimpanzee iPSCs injected into 99 macaque 5-day-old embryos formed cross-species chimeras two days after injection. Strikingly, the chimpanzee iPSCs survived, proliferated and integrated near the inner cell mass (ICM) of rhesus macaque embryos. These findings highlight the broad potential of primate iPSCs in forming cross-species chimeras beyond rodents and provides a foundational basis for organ generation using human iPSCs.

## Introduction

Induced pluripotent stem cells (iPSCs) have the potential to develop into all three germ layers of ectoderm, mesoderm and endoderm and form functioning organs *in vivo*. A robust validation of iPSC pluripotency is evaluating whether these cells can survive and propagate along with the endogenous embryonic cells when injected into a developing blastocyst of the same species or a closely related species (*1*). The pluripotency of mouse iPSCs has been successfully demonstrated by injection into both mouse and rat blastocysts, after which the cells survived and formed mouse-mouse or mouse-rat cross-species chimeras respectively (*2*). These cross-species rodent chimeras were further used to grow mouse organs (e.g., pancreas) in rats (*Pdx1*−/− apancreatic rat) and vice versa (*2, 3*). In principle, this approach could be applied to generate human organs in cross-species chimeras. Although some human or primate iPSCs, when injected into mouse blastocysts, can occasionally contribute to low-level chimerism in early mouse embryonic stages (*4–7*), further development of human-mouse chimeras is challenging likely due to long evolutionary divergence and developmental differences. A better model is therefore needed to validate human iPSC potency and explore *in vivo* generation of human organs.

Compared to rodents, non-human primates are a more translationally relevant species to corroborate cross-species *in vivo* organogenesis, however several issues preclude this approach. Ethical concerns restrict injecting human iPSCs into non-human primate (NHP) embryos due to the unknown consequences of chimeric brain and gamete contribution(*8, 9*). Technical issues also remain. For example, only PSCs in the naïve state (e.g., mouse, rat) can survive and form chimeras if injected into blastocysts. In contrast, mouse epiblast-derived stem cells (EpiSCs) are in the primed state of pluripotency and fail to survive and form chimeras in rodent blastocysts. The common cell-culture methods for primate iPSCs results in primed-like characteristics and are not expected to be chimera competent(*10*). All efforts so far have been in the context of within-species chimerism (*11–13*) and it is not yet clear whether primate iPSCs can survive and form cross-species embryos when injected into early-stage embryos (e.g., blastocyst) of closely-related primate species.

In this study, we explored the ability for cross-species NHP iPSCs to engraft rhesus macaque blastocysts. Pig-tailed macaque and chimpanzee were chosen as the iPSC donor-species since their evolutionary divergence from rhesus macaque is similar in length to successfully generated cross-rodent chimeras. Importantly, as the closest relative to humans, chimpanzee-derived iPSCs may serve as an ethically-mindful surrogate for human iPSCs when exploring primate pluripotency and chimeras. Therefore, we derived pig-tailed macaque and chimpanzee iPSC lines and optimized culture conditions. Similar functional and developmental characteristics of chimp and human iPSCs was confirmed through co-culture experiments *in vitro.* We then injected the NHP iPSCs into rhesus macaque embryos. High *BCL2* expression was necessary to overcome primed-stage challenges and enable engraftment. Both chimpanzee and pig-tailed macaque iPSCs survived and proliferated near the inner cell mass (ICM) in the rhesus macaque blastocyst two days after iPSC injection. Together, these results are the first steps toward forming cross-species chimeras using iPSCs between the species of primate taxa. These results suggest that *BCL2*-expressing primate iPSCs might be a valuable tool for the formation of cross-species chimera for ultimate goal of organ generation.

## Results

### NHP iPSC generation, culture condition optimization, and validation

We derived iPSCs from peripheral blood mononuclear cells (PBMCs) of two NHP species, chimpanzee (*Pan troglodytes*) and pig-tailed macaque (*Macaca nemestrina*), using CytoTune kit (Sendai virus containing Oct3/4, Sox2, c-Myc, and Klf4) (*14*) (Supplementary figure S1B). We optimized NHP iPSC culture conditions by using the pluripotent stem cell culture medium E8 (*15, 16*), supplementing with 2μM XAV939 (“E8X”) to inhibit Wnt/β-catenin canonical pathway. Compared to E8 medium, the use of E8X reduced spontaneous differentiation of NHP iPSCs (Supplementary figure S1D). We also tested if feeder-free conditions using iMatrix-511 would be sufficient for maintenance of pig-tailed macaque iPSC (data not shown). Unlike chimpanzee iPSCs which exhibited little spontaneous differentiation when seeded on plates coated with iMatrix-511 (Supplementary figure S1D), pig-tailed macaque iPSCs did differentiate and presented the best morphology and the least spontaneous differentiation when cultured on mouse embryonic fibroblasts (MEF) in E8X medium (Supplementary Figure S1D). In addition, XAV939 was critical for long-term maintenance of pig-tailed macaque iPSC on MEFs (Figure 1A). We also evaluated the expression of TRA-1-60, a pluripotency marker (*17, 18*), in NHP iPSCs in E8 and E8X medium. We observed that the proportion of TRA-1-60 expressing cells significantly decreased after one passage in E8 medium (Supplementary Figure S2). We further validated the normal karyotype of the generated iPSC lines for both species (Supplementary Figure S1C), in vitro differentiation potential to cranial neural crest (Supplementary Figure S4) and the ability to form teratomas (“teratoma assay”) in SCID mice (Figure 1D and Supplementary Figure S3). These assays confirmed genomic integrity and pluripotency features of both the chimpanzee and pig-tailed macaque iPSCs.

**Figure 1.**
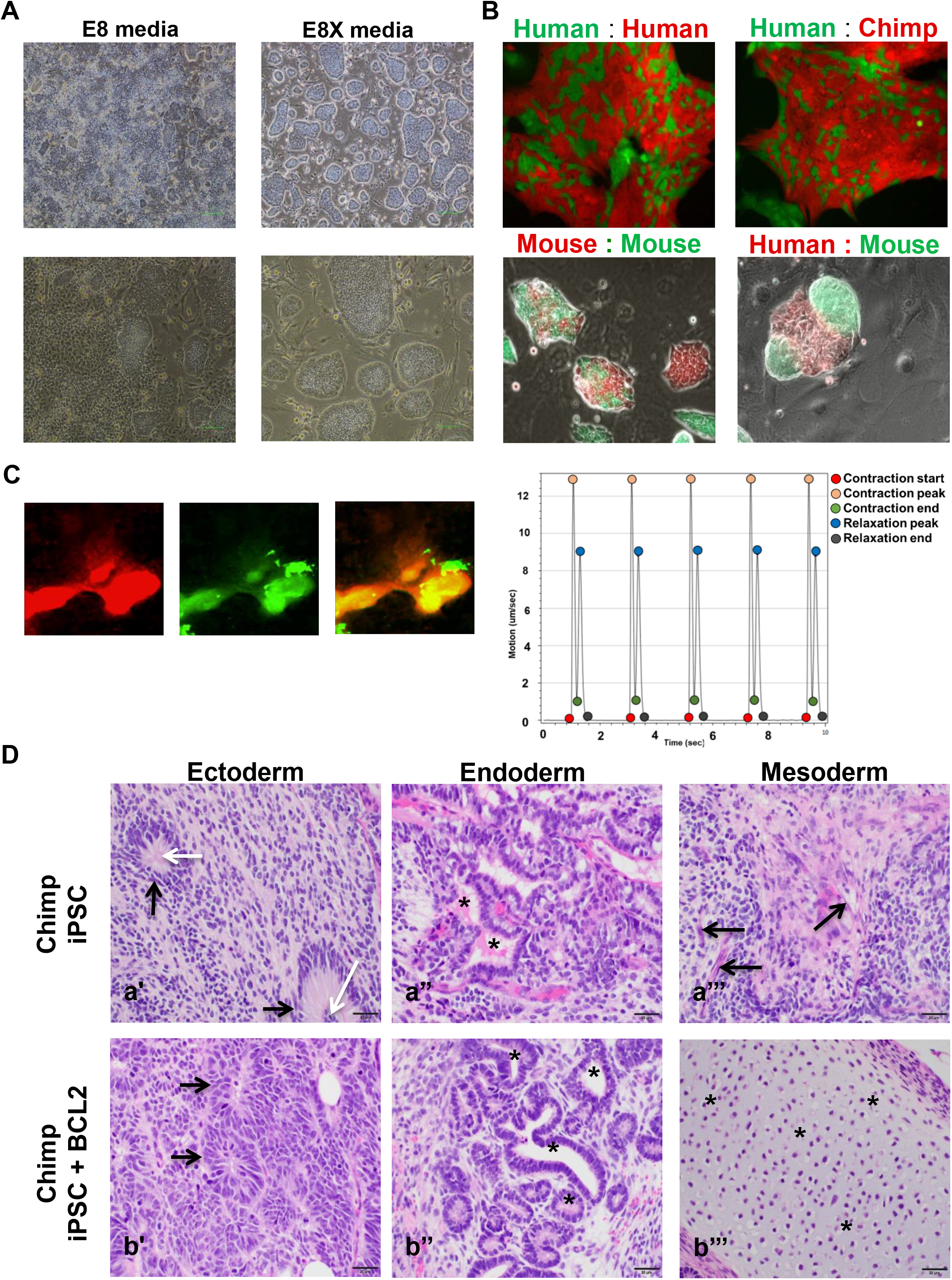
NHP iPSC media optimization and co-culture. A) Colony morphology of pig-tailed macaque iPSC in NHP-E8X media on MEF at 4X and 10X resolution (left) and the colony morphology of pig-tailed macaque iPSCs in E8 media on MEF at 4X and 10X resolution (right). B) Co-culture of human-human iPSC (top-left) and human-chimp iPSC (top-right). Human and chimpanzee iPSCs mix and integrate homogenously without forming segregated colonies. Co-culture of mouse-mouse iPSCs form mixed and homogenously integrated colonies (bottom-left). Human and mouse iPSCs form separate and segregated colonies (bottom-right). C) Cardiomyocyte layers differentiated from the co-culture mixture of chimpanzee and human iPSCs. The tdTomato-fluorescent labeled beating chimpanzee iPSC (red, left) and the human GFP-fluorescent labeled beating cardiomyocyte (green, middle) and the overlay image of the mixed cardiomyocytes from the mixture of chimpanzee and human iPSC (Supplementary video 1). Electrocardiogram of cardiomyocyte derived from the mixture of human and chimpanzee iPSCs indicating contraction and relaxation start, peak, and end (right). D) Hematoxylin and eosin (H&E)-stained sections of chimpanzee teratoma showing all three embryonic germ layers, derived from chimpanzee iPSCs with and without Bcl2 transduction. Ectoderm (a’, b’) is characterized by the presence of neural ectoderm arranged in rosette-like patterns (black arrows, a’ and b’), that resemble embryonic neural tubes (white arrows, a’). Endoderm (a’’, b’’) is characterized by the presence of glandular and/or secretory epithelium (black asterisks). Mesoderm (a’’’, b’’’) is characterized by the presence of elongate, mesenchymal cells (black arrows, a’’’) and cells embedded in a pale blue gray hyaline matrix resembling articular cartilage (black asterisk, b’’’). Magnification: 40x. Scale bars: 20μm.

### *In vitro* co-culture of NHP and human iPSC

We next tested functional and developmental similarities of chimpanzee and human iPSCs in an *in vitro* culture setting by co-culturing human and chimpanzee iPSCs. We mixed equal number of single-cell dissociated human and chimpanzee iPSCs and seeded ~40,000 cells (20,000 human iPSCs and 20,000 chimpanzee iPSCs) on plates coated with iMatrix-511. Co-culturing human and chimpanzee iPSCs indicated that the two cell types do not form distinct and separate iPSC colonies. Instead, the iPSCs mixed homogenously in a blended manner similar to that of co-culturing two different human iPSC cell lines (Figure 2B). In contrast, co-culture of human iPSCs and mouse EpiSCs resulted in separate distinct colonies (Figure 2B). Thus, chimpanzee and human iPSCs appear functionally similar *in vitro*, in contrast with human and rodent iPSCs. We then differentiated the mixture of human and chimpanzee iPSCs into cardiomyocytes. We observed the mixture of human and chimpanzee differentiating iPSCs resulted in beating cardiomyocyte aggregates composed of cells from both human and chimpanzee (Figure 1C, Supplementary video 1 and 2). Mixed beating cardiomyocyte colonies were not obtained using human iPSCs mixed with either mouse EpiSC or pig-tailed macaque iPSCs. Based on these experiments, we conclude that chimpanzee iPSCs behave very similarly to human iPSCs during *in vitro* culture and differentiation. Thus, although the use of human iPSCs for embryonic cross-species chimera studies is restricted, these results suggest that chimpanzee iPSCs maybe functionally equivalent to human iPSCs.

**Figure 2.**
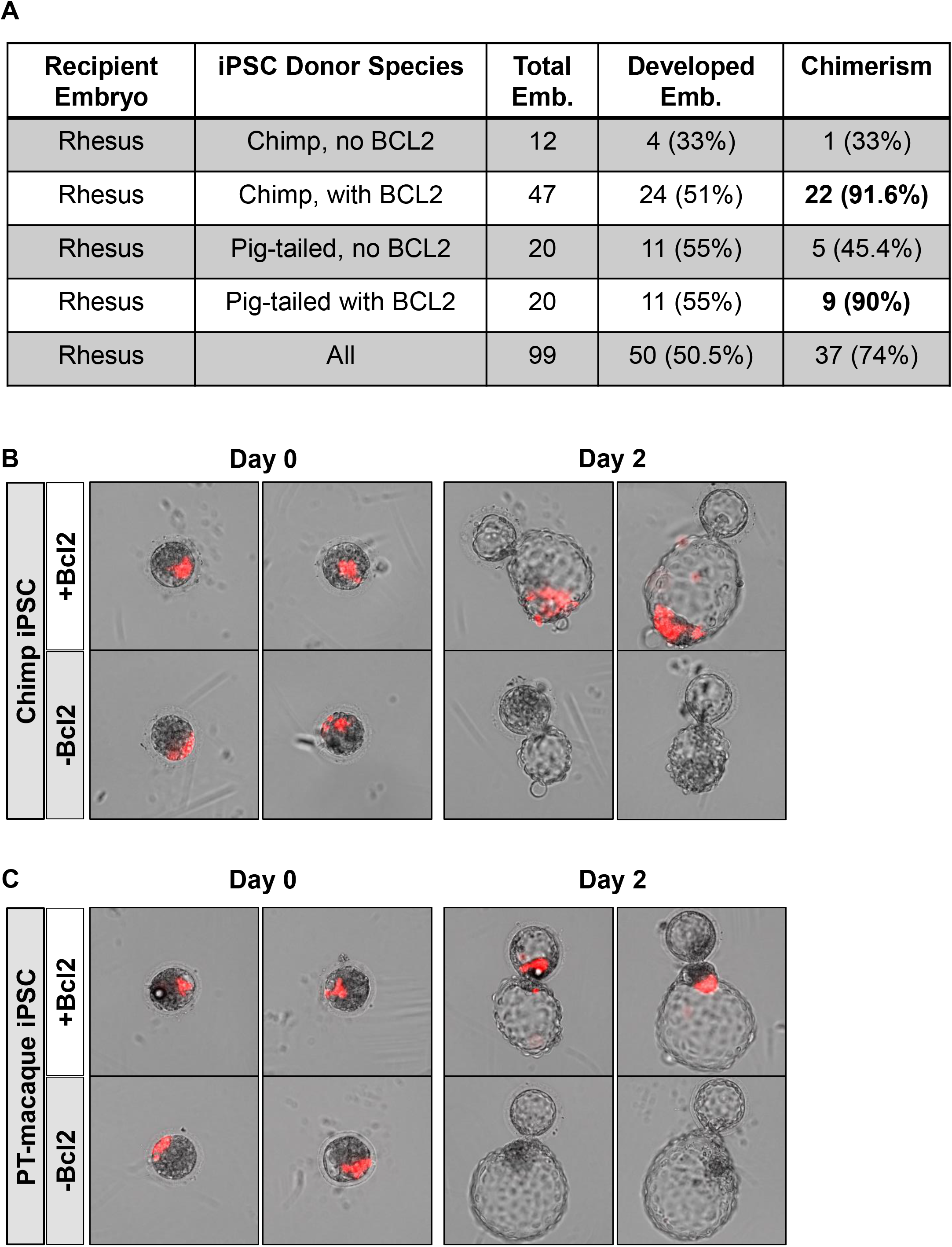
BCL2 expression supports *In vitro* primate cross-species chimerism. (A) The summary of the number of both chimp and pig-tailed with or without Bcl2 into rhesus. Red fluorescent BCL2-expressing (B) chimpanzee or (C) pig-tailed macaque iPSCs survive, proliferate and localize near rhesus ICM 2 days post injection imaged through in vitro culture.

### Assessment of *BCL2*-transduced human, chimp and pig-tailed macaque iPSC cells in a mouse teratoma model

To overcome apoptosis associated with primed-state iPSC injection into blastocysts, we expressed the anti-apoptotic gene *BCL2* in the iPSCs. We then evaluated whether *BCL2* expression in human, chimpanzee, and pig-tailed macaque iPSCs affected the differentiation potential of these cells. We first tested whether these primate *BCL2*-expressing iPSCs form all three germ layers (ectoderm, mesoderm and endoderm) when injected into NOD mice similar to non-BCL2 transduced iPSCs (see methods for the assays). Teratomas developed from chimpanzee iPSCs and pig-tailed macaque iPSCs with and without *BCL2* in the testes and subcutaneous tissues of NOD mice. The changes in the size of the tumor in each mouse was recorded every 3 days and an estimate of the tumor size was made over time. At the end of the teratoma study on day 62, tissues from the NOD mice were thoroughly examined histologically. Tissues examined include dorsal haired skin, reproductive tract, liver, gallbladder, spleen, kidneys, adrenal glands, pancreas, salivary gland, mandibular lymph nodes, heart, lung, trachea, esophagus, tongue, eyes, brain, and cerebellum. There was no evidence of metastasis nor any other abnormality in any of the examined tissues. Differentiated cells were observed regardless of *BCL2* transduction (Figure 1D and Supplementary Figure S3A). Similar results were found using the human H9 cell line and *BCL2*-expressing H9 cell line (Supplementary Figure S3B). These results indicate that expression of *BCL2* in primate pluripotent stem cells does not negatively affect the differentiation potential of these cells.

### *In vitro* evaluation of nonhuman primate cross-species chimerism

To test whether chimpanzee and pigtail macaque cells can successfully inhabit early embryos of rhesus macaques, we injected iPSCs into rhesus macaque embryos. Follicle retrieval from rhesus macaques was performed by ultrasound-guided aspiration of follicles that occurred roughly 33 hours following an hCG administration. Oocyte insemination and embryo culture were performed according to our standard in vitro protocols (see methods for the details) (*19, 20*). 8-12 chimpanzee or pig-tailed macaque red fluorescent tagged iPSCs were injected into 99 rhesus embryos (59 with chimpanzee and 40 for pig-tailed macaque iPSCs). Two control embryos underwent injection without iPSCs. Two days after injection, 22 out of 24 (91.6%,) of the rhesus embryos injected with chimpanzee *BCL2*-expressing iPSCs survived and chimpanzee iPSCs proliferated inside the rhesus macaque embryos near the ICM, as indicated by expanded red florescent areas in the embryos (Figure 2B). Similar results were observed after injection the *BCL2-*expressing PT-macaque iPSCs (9 out of 11 (90%), Figure 2C). In contrast, most of the non-*BCL2*-expressing iPSC (33% chimpanzee and 45.5% pig-tailed macaque) did not survive in rhesus blastocysts during this time period (Figure 2A). Overall, approximately 50.5% of the injected embryos survived and developed in vitro after injection of iPSCs. This is consistent with the normal survival developmental rate of rhesus macaque blastocysts. These results indicate that the injected cells survive, proliferate and integrate into ICM.

## Discussion

Our results indicate that the use of E8X culture medium improved the maintenance of NHP iPSCs. In chimpanzee iPSCs, the addition of XAV939 reduces spontaneous differentiation; however, in pig-tailed macaque iPSCs, addition of XAV939 is crucial for maintenance of iPSCs, as the majority of iPSCs without the XAV939 in their medium differentiate (Figure 1A). The inhibition of Wnt/β-catenin in N2B27 has been shown to improve maintenance of mouse EpiSCs and our results indicate that addition of XAV939 in E8 medium helps maintain pluripotency in NHP iPSCs (*21, 22*).

We also show that *BCL2*-transduced chimpanzee and pig-tailed macaque iPSCs can survive, proliferate, and aggregate near rhesus macaque ICMs for at least two days after blastocyst injection. These results are the first promising steps to evaluate whether primate iPSCs can form cross-species chimeras. It was previously shown that mouse EpiSCs, which resemble the primed state of pluripotency and are not normally chimera competent, were only able to engraft rodent blastocysts with transient expression of *BCL2* (*23*). In agreement with those results, minimal chimerism was observed following blastocyst injection of non-*BCL2*-expressing chimpanzee and pig-tailed macaque iPSCs. Importantly, the NHP *BCL2*-expressing iPSCs aggregate in or around the ICM, which is the region of the blastocyst that develops into the embryo proper. Therefore, these findings confirm the potential of *BCL2*-expressing primate iPSCs to form chimeras in a different species of nonhuman primates (e.g., rhesus macaque). This promising result among NHP species is consistent with previous findings regarding the chimera-forming potential of *BCL2*-transduced mouse and human cells injected into mouse embryos (*23, 24*). Notably, the evolutionary divergence of chimpanzee and rhesus macaque is similar to that of rat and mouse (~20 million years), which has been the most successfully developed model for mammalian cross-species chimeras and *in vivo* organogenesis.

Due to the ethical considerations regarding injection of human iPSCs into nonhuman primate embryos (*1, 8*), we did not inject human iPSCs into the rhesus blastocysts. Instead, we injected chimpanzee iPSCs, the closest relative to humans (~6 million years). We showed the compatibility of the chimpanzee and human iPSCs by co-culturing and co-differentiating the iPSC mixture into beating cardiomyocyte nodes. This indicates that, at least *in vitro*, human and chimp cells respond to similar development cues and can form homogenously mixed structures. Thus, cross-species experiments utilizing chimpanzee iPSCs could likely be extrapolated to human iPSCs.

*BCL2*-transduced NHP iPSC did not result in any abnormal histopathology in mice when injected into the testes or subcutaneous tissues to form teratomas. From these results, we hypothesize that *BCL2*-transduced NHP iPSCs will develop into three distinct germ layers and form NHP cross-species chimeras without significant histologic alteration in a NHP chimeric fetuses. In this study, we focused only on the potential of *BCL2*-transduced cells in forming early NHP cross-species chimeras *in vitro*.

Formation of chimeras using human iPSCs with large animals (e.g., pigs) has been difficult (*25*). Other models are needed to explore chimera formation and cross-species organ generation in a practical and ethically safe manner. Our results indicate that chimpanzee-rhesus macaque chimeras are likely to be more successful and relevant to human organogenesis. To the best our knowledge, these are the first positive results indicating that chimpanzee and pig-tailed macaque iPSCs can survive and proliferate in rhesus macaque early embryos. The next step is to conduct embryo transfer of the chimeric NHP embryos into a rhesus macaque female. This will determine whether *BCL2*-expressing chimp iPSCs can form a cross-species chimera *in vivo*, investigating the potential of cross-species chimera formation among nonhuman primate species. These studies using chimpanzee iPSCs will serve as a useful model for future studies of human organ generation.

## Author contribution

MR conceived the study, species selection, iPSC derivation, and NHP iPSC media optimization. KL guided on iPSC medium optimization. MR and FS conducted iPSC fluorescent-labeling, single cell expansion and BCL2 transduction. CAV conducted rhesus macaque oocyte retrieval, IVF, and embryo generation. FS conducted embryo injection and imaging. MR, FS, JB conducted iPSC co-culture experiments. MR, VKB, and AO conducted cardiomyocyte differentiation and evaluation. VKB conduced ectoderm differentiation. MR and JGV conducted BCL2 teratoma assays and histopathology analysis. JCW supervised cardiomyocyte experiments. CAV, HN, and MPS supervised the study.

## Acknowledgment

We would like to thank David Glenn Smith for consulting on NHP species selection and evolutionary distance among NHP species that apply to pluripotent stem cell biology.

---

**Supplemental Figure S1.**
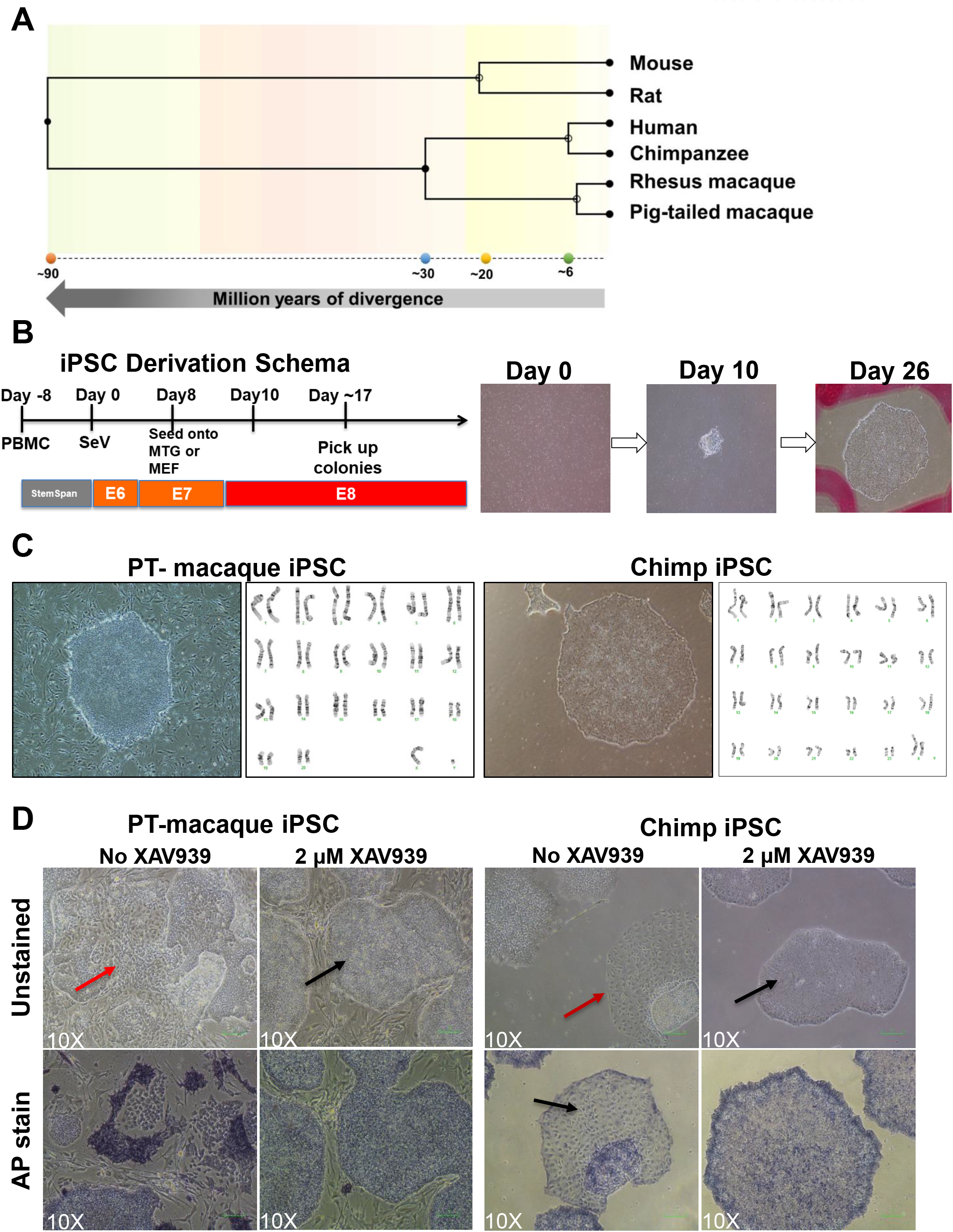
A) Schematic figure of evolutionary dstance among the primate species of the study. B) Schematic figure of reprogramming timeline and media used for reprogramming chimpanzee and pig-tailed macaque PBMCs into iPSCs (left) and the morphology of chimpanzee reprogramming cells into iPSCs at days of 0, 10 and 26 of reprogramming. C) Colony morphology and the normal karyotype of the pig-tailed macaque iPSCs (left) and chimpanzee iPSCs (right). D) Morphology of pig-tailed macaque iPSCs on MEF with and without XAV939 in the E8 media (left) and chimpanzee colonies on iMatrix with and without XAV939 in the E8 media (right). Black arrows indicate the differentiating cells surrounding the iPSC colonies in the media without XAV939 and red arrows indicate normal iPSC morphology in E8 media with XAV939.

**Supplementary Figure S2.**
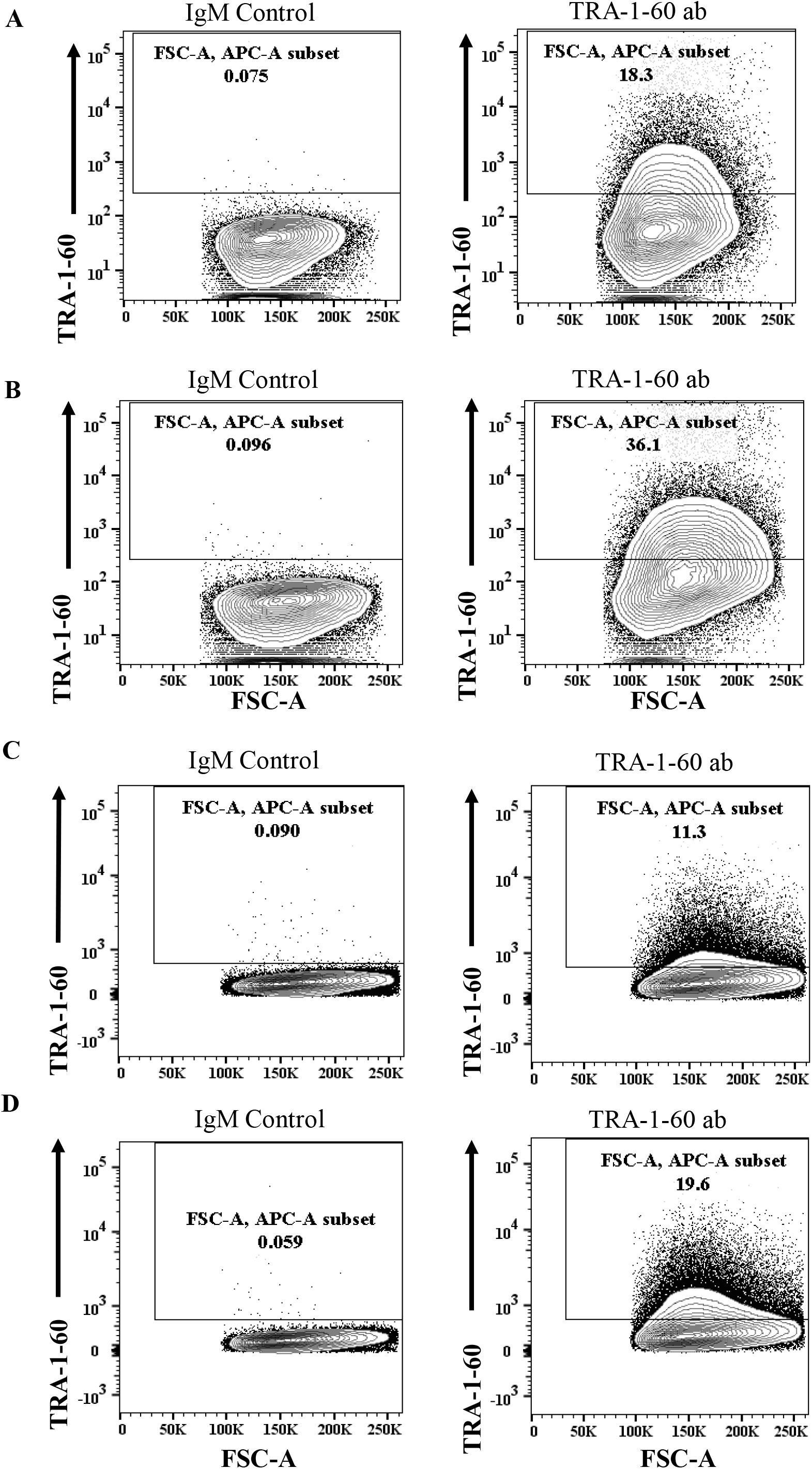
Expression of pluripotency marker TRA-1-60 in pig-tailed macaque and chimpanzee iPSCs. (A) Flow cytometry results of TRA-1-60 expression in pig-tailed macaque iPSCs in E8 media. The IgM control antibody (left) and the TRA-1-60 stained antibody indicating 18.3% of the parental cells in E8 medium express TRA-1-60 (right). (B) Flow cytometry results of expression TRA-1-60 in pig-tailed macaque iPSCs in E8X media. The IgM control antibody (left) and the TRA-1-60 stained antibody indicating 36.1% of the parental cells express TRA-1-60 (right). (C) Flow cytometry results of TRA-1-60 expression in chimpanzee iPSCs in E8 media. The IgM control antibody (left) and the TRA-1-60 stained antibody indicating 11.3% of the parental cells in E8 medium express TRA-1-60 (right). (D) Flow cytometry results of TRA-1-60 expression in chimpanzee iPSCs in E8X media. The IgM control antibody (left) and the TRA-1-60 stained antibody indicating 19.6% of the parental cells express TRA-1-60 (right).

**Supplemental Figure S3.**
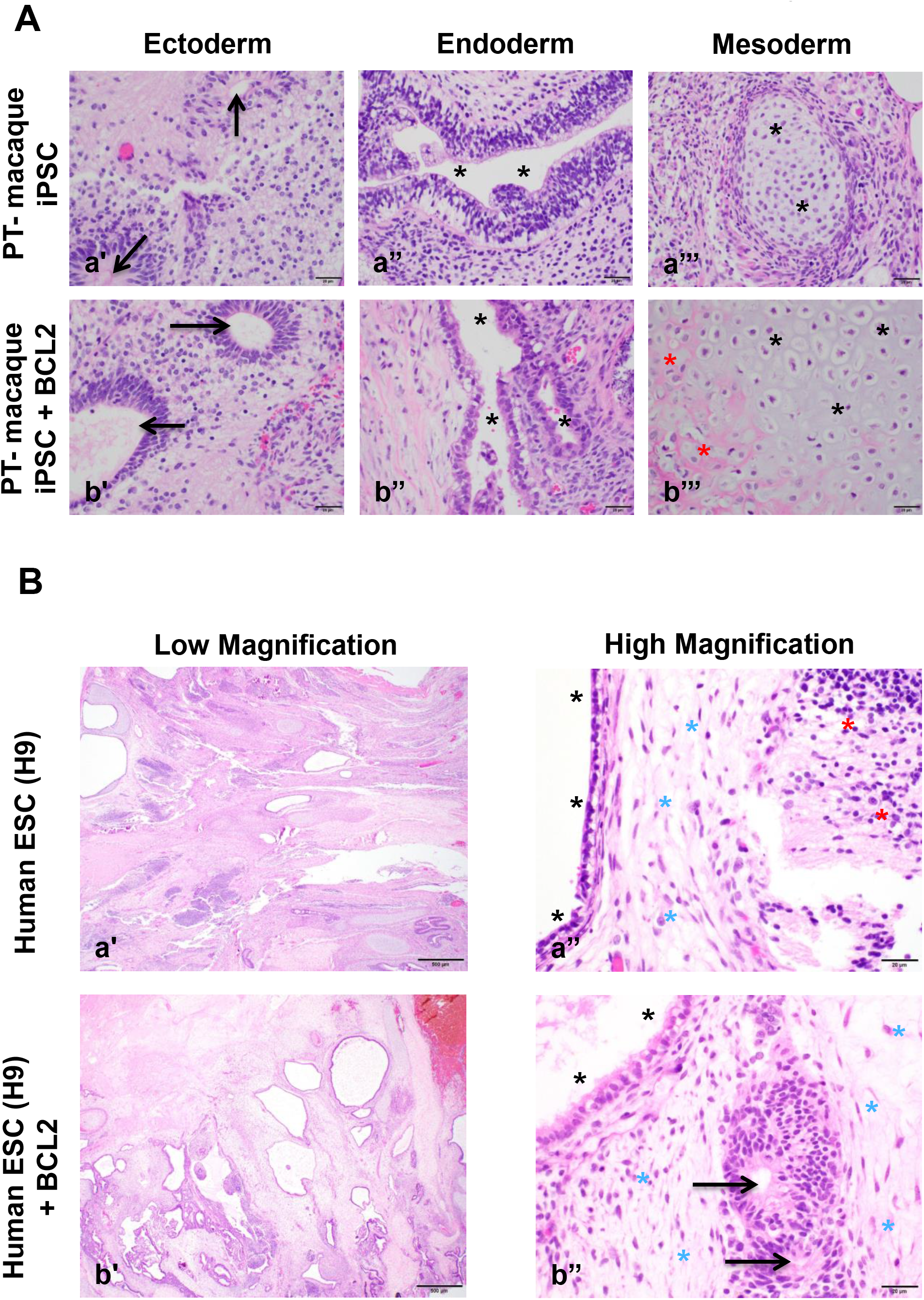
Histopathology of teratoma from iPSCs with and without BCL2. (A) Hematoxylin and eosin (H&E)-stained sections of pig-tailed macaque iPSC teratoma showing all three embryonic germ layers, derived from pig-tailed macaque iPSCs with and without Bcl2 transduction. Ectoderm (a’, b’) is characterized by the presence of neural ectoderm arranged in rosette-like patterns that sometimes resemble embryonic neural tubes (black arrows, a’ and b’). Endoderm (a’’, b’’) is characterized by the presence of glandular and/or secretory epithelium (black asterisks). Mesoderm (a’’’, b’’’) is characterized by the presence of cells embedded in a pale blue gray hyaline matrix resembling articular cartilage (black asterisks, a’’’ and b’’’), or in a pale eosinophilic matrix resembling bone (red asterisks, b’’’). (B) Hematoxylin and eosin (H&E)-stained sections of teratoma showing all three embryonic germ layers, derived from H9 (human ESCs) cell line with and without Bcl2 transduction. Similar to the teratomas derived from chimp and pig-tailed macaque iPSCs, these tumors have all three embryonic germ layers represented, including ectoderm, endoderm, and mesoderm (a’-b’’). Ectoderm is characterized by the presence of tissue resembling neuropil (red asterisks, a’), or rosette-like arrangements resembling neural tubes (black arrows, b’’). Endoderm is characterized by the presence of glandular and/or secretory epithelium (black asterisks, a’’ and b’’). Mesoderm is characterized by the presence of elongate cells in a loose matrix resembling embryonic mesenchyme (blue asterisks, a’’ and b’’). Magnification: a’, b’= 2x; a’’, b’’= 40x. Scale bars: a’, b’= 500 μm; a’’, b’’= 20 μm.

**Supplementary Figure S4.**
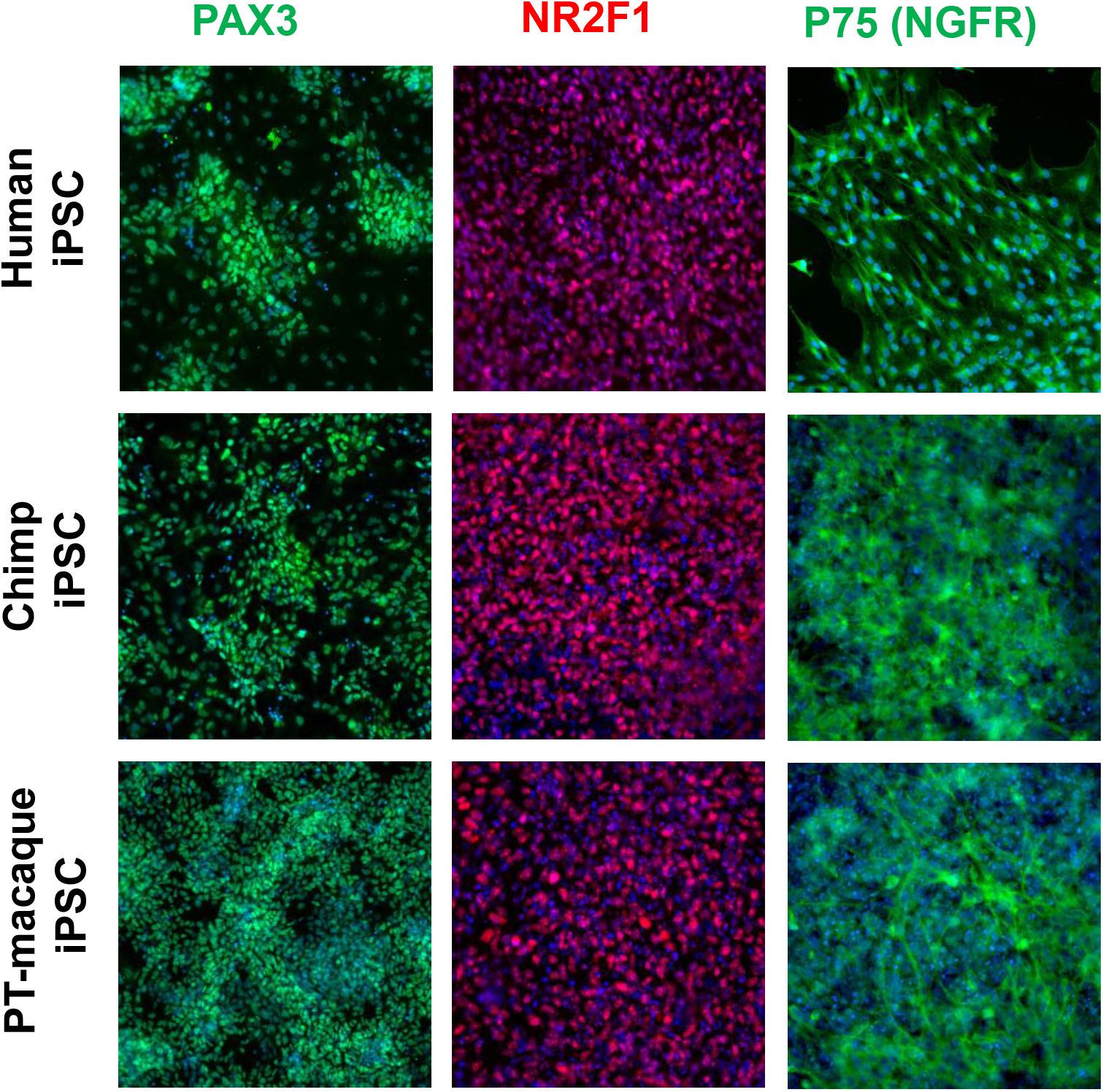
Differentiation of pig-tailed macaque, chimpanzee, and human iPSCs into cranial neural crest.

**Supplementary Figure S5.**
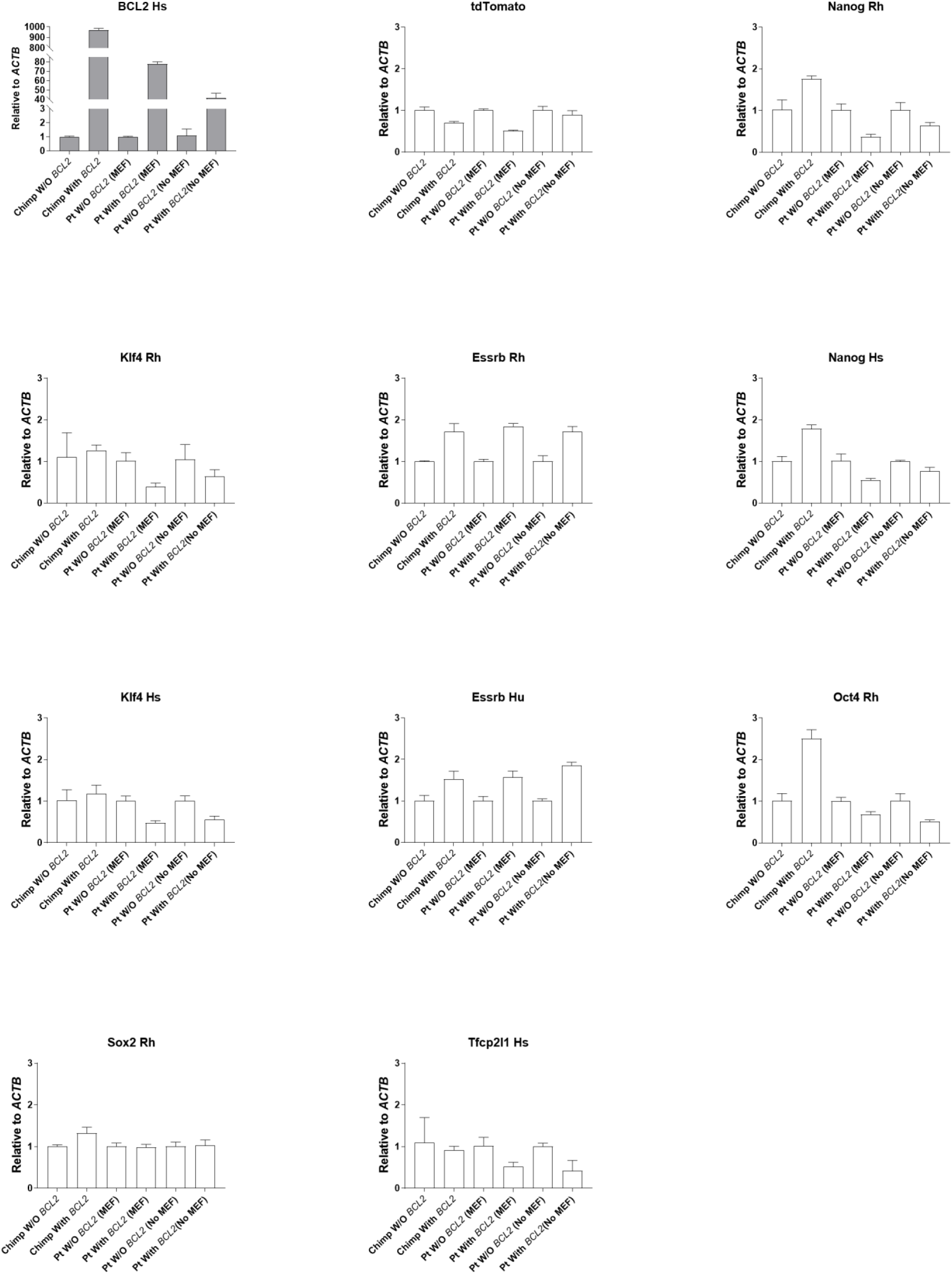
Expression of *BCL2* and pluripotency markers in iPSCs from chimpanzee and pig-tailed macaque in with and without *BCL2* transduction. Different qPCR probes from both human (Hs) and rhesus macaque (Rh) were used for the pluripotency marker genes.

## Methods

**All the experiments in this study were conducted following the Institutional Animal Care and Use Committee (IACUC) protocol at University of California Davis, and Administrative Panel on Laboratory Animal Care (APLAC) protocol at Stanford University, Stem Cell Research Oversight (SCRO) at Stanford University and Administrative Panel on Biosafety (APB) protocol at Stanford University.**

### iPSCs derivation and maintenance

a) Chimpanzee (*Pan troglodytes*) iPSCs: chimpanzee peripheral blood mononuclear cells (PMBCs) were obtained from Southwest National Primate Research Center (SNPRC), San Antonio, Texas, USA according to the corresponding SNPRC IACUC protocol. and the PBMCs were reprogrammed into induced pluripotent stem cells (iPSCs) optimizing a feeder free condition(*26*). In summary, PBMCs were cultured to expand blood progenitor cells in Stem Span medium (Stemcell Technologies Inc. Vancouver, Canada) for 9 days. The next day (day of reprogramming, “day 0”), the cells were transfected with Sendai virus with four transcription factors including c-myc, KLf4, Sox2, Oct3/4 using CytoTune 2.0 kit (ThermoFisher Scientific, Waltham, MA, USA). On the day after transfection (day 1), all the cells transfected with Sendai virus were transferred into the plates coated with Matrigel Matrix (Corning Inc., Corning, NY, USA) in StemSpan medium (Stemcell Technologies Inc. Vancouver, Canada). On day 2, floating cells were transferred to new wells of a Matrigel-coated 6-well plate in 2 mL of Stem Span medium. On day 3, 2ml of Essential-6 medium (E6) (ThermoFisher Scientific, Waltham, MA, USA) were added into existing 2mL StemSpan medium and cells in the 6-well plates, and the mixture was left unchanged on day 4. On day 5, the media in each well was removed and replaced by 2mL E6 medium, and left unchanged on day 6. On day 7, all medium was removed and 2mL of E6 medium + 100 ng/mL of bFGF (ThermoFisher Scientific, Waltham, MA, USA) were added. On days 8, 9, and 10 the medium was replaced with 2mL E7 (Essential 6 medium + 100ng/ul of FGF). Starting on day 11, the medium was replaced by E8 (*27*)until days 16 to day 18 when the first colonies of iPSCs were observed. Each colony was manually picked with 1000μL pipet and was cultured on new Matrigel-coated plates in E8 medium with 2nM of Thiozivine (TZV) on the first day after passage. After the first passage, 2nM of Wnt inhibitor (XAV939) were added to the E8 medium which resulted in colonies with little-to-no differentiation. For the subsequent passaging, the wells were coated with 5μL of iMatrix-511 (Takara Bio Inc. Kusatsu, Japan) in 2mL of the E8 + E8 specific supplement + XAV939 (2nM) for expansion.

b) Pig-tailed macaque (*Macaca nemestrina*) iPSC: Blood samples were obtained from a 12-year old male pig-tailed macaque from the Johns Hopkins University colony. The PBMC (peripheral blood mononuclear cells) were isolated from whole blood in each sample and were then reprogrammed into induced pluripotent stem cells (iPSCs) making modifications and optimizing a previously published feeder free protocol(*26*). In summary, PBMCs were cultured in medium to expand blood progenitor cells in StemSpan medium for 9 days. The next day, the cells were transfected with Sendai virus of four transcription factors c-myc, KLf4, Sox2, Oct3/4 using CytoTune 2.0 kit (ThermoFisher, Waltham, MA, USA) on the day of reprogramming (day 0). On the day after transfection (day 1), all the cells transfected with Sendai virus were transferred into a 6-well plate (30,000 cells per well) each well containing ~150,000 mouse embryonic fibroblast (MEF) feeder cells (Stemcell Technologies Inc. Vancouver, Canada). The MEF plate preparation was conducted the day before plating the reprogramming cells as explained elsewhere (*28*). On day 2, the floating cells were transferred to new wells of 6-well plate containing MEF in 2 mL of the Stem Span media. On day 3, 2mL of in Essential-6 media (ThermoFisher Scientific, Waltham, MA, USA) were added into existing 2mL StemSpan medium on MEF in which transfected cells were. No medium change on day 4 was conducted. On day 5, all the medium in each well were removed and replaced by E6 medium. No medium change on day 6 was conducted. On day 7, all medium was removed and 2mL of E6 + 100 ng/mL of bFGF (Stemcell Technologies Inc. Vancouver, Canada) were added. On day 8, 9, and 10 media was changed and with E7 (E6 + 100ng/ul of FGF). From day 11, the media was replaced by E8 until days 16 to day 18 when the first colonies of iPSC were observed. Each colony was manually picked with 1000μL pipet and were cultured on new matrigel-coated plated in E8 media and 2nM of TZV on the first day after passage. After the first passage, 2nM of Wnt inhibitor (XAV939) were added into the E8 media which resulted in colonies with little-to-no differentiation. For subsequent passaging and expansion, the iPSCs were washed with PBS and were incubated at using 1mL of Accumax (Stemcell Technologies Inc. Vancouver, Canada) for one well of a 6-well plate for 5 minutes at 37°C. The Cells were then aspirated, and single cell dissociated using a 1000μL. The cells were suspended in E8X medium with 2nM of TZV on mouse embryonic fibroblast cells (MEF) for each passaging during the expansion and maintenance.

#### iPSCs Karyotyping

The macaque-derived (*Macaca nemestrina*) cell lines (Ma2E8) was harvested by standard cytogenetic methodology of mitotic arrest, hypotonic shock and fixation with 3:1 methanol-acetic acid. Chromosome slide preparations were stained by G-banding and classified by the standard *M. nemestrina* karyotype 1, 2. Analysis of 20 metaphase cells demonstrated an apparently normal male karyotype of 20 autosomes and two sex chromosome (XY) in all cells (*29, 30*).

#### iPSCs labeling with fluorescent vector and Bcl2 vector

The iPSC from chimpanzee and pig-tailed macaque were transfected with the tdTomato (tdT) red fluorescent vector (*23*). The tdT transfected cells were then clonally expanded from single cells to make sure all the cells from each clone contained the same number of tdTgenes. This was done to minimize the heterogeneity in expression of tdTomato among iPSC. The tdTomato single cells expanded iPSC were subsequently transfected with the Bcl2 vector containing neomycin resistance gene(*23*). The iPSCs then were expanded clonally from single cells and treated with neomycin for two weeks to obtain iPSCs with same number of integrated BCL2 in each cell. The iPSCs containing tdT and Bcl2 were then used for the injection into rhesus macaque embryos.

#### Alkaline Phosphatase (AP) Stain

Cells were rinsed twice with DPBS and fixed for five minutes at room temperature (RT) with 4% PFA. Then 1-Step™ NBT/BCIP Substrate Solution (Cat No. #34042, ThermoFisher Scientific, USA) was added to the wells and allowed to develop for 8-10minutes at RT. The solution was aspirated and the wells were washed twice with DPBS before being imaged.

#### Co-culture experiment of iPSCs

Equal numbers iPSCs from each species was used for the paired species co-culture experiment. The paired iPSCs from human and chimpanzee, human and human, human and mouse, and mouse and mouse (40,000 cells from each species) were cultured to perform co-culture experiments. The differentiation into cardiomyocytes was conducted on these co-cultured cells as described in iPSCs into cardiomyocyte differentiation section (below).

#### Teratoma assay

The iPSCs were cultured and dissociated using Accumax (Stemcell technologies Inc. Vancouver, Canada) in E8 + supplement and XAV939 (2nM) and TZV (2nM). The cells were washed once with E8 + supplement and XAV939 (2nM) and TZV (2nM) media (75%) and 25% DMEM-F12 in total concentration 70 million cells /mL. 30μL of the solution containing total of 2.1 million cells (or placebo controls) were injected subcutaneously on the dorsum of each mouse. The same number of cells were injected into the testes. Tumors were monitored and tumor size was measured, every other day for 60 days.

#### Histopathology

At the end of the study, mice were humanely euthanized via CO_2_ inhalation. Tissues were assessed for gross evidence of tumor burden, and tissues were collected in 10% neutral buffered formalin (NBF). After fixation, tissues were routinely processed, paraffin embedded, sectioned at 5.0μm, and sections routinely stained with hematoxylin and eosin (H&E). Tissue sections were examined for the presence of teratomas. Teratomas were defined as tumors of embryonic origin containing the three embryonic germ layers (endoderm, mesoderm, ectoderm) and/or their derivatives. When present, teratomas were assessed for evidence of malignant transformation such as local invasion into the adjacent tissues. Histologic assessment was performed using an Olympus BX43 upright brightfield microscope, and images were captured using the Olympus cellSens software.

#### qPCR

qPCR for pluripotency markers were conducted using the conserved probes designed to work for a wide range of primate species including both human and rhesus macaque. A list of all pluripotency genes measured via qPCR and the corresponding primers and probes are listed in supplementary table (Supplementary Table 1)

### Rhesus macaque oocyte retrieval, fertilization and embryo culture

The Controlled Ovarian Stimulation (COS) protocol overrides follicle selection, resulting in numerous preovulatory follicles that respond to a timed injection of hCG to simulate the natural LH surge. One adult female rhesus monkey was used for oocyte retrieval and was housed at the California National Primate Research Center (CNPRC) in accordance with the ethics guidelines established and approved by the Institutional Animal Use and Care Administrative Advisory Committee at the University of California-Davis (*31*). The COS cycle began within four days of menses as described previously with twice-daily injections of human recombinant FSH (37.5 IU) for seven days followed by a single hCG injection (1,000 IU) (*32, 33*). Follicle retrieval was performed by ultrasound-guided aspiration of follicles that occurred approximately 33 hours following hCG administration. Oocyte insemination and embryo culture were performed according to our standard in vitro protocols(*19, 20*). Approximately 5 days after insemination, embryos were transported from CNPRC to Stanford using a portable incubator that has been shown to maintain embryo viability(*34*).

### Injection of NHP iPSC into the rhesus macaque embryos and imaging

In preparation for microinjection, the pig-tailed macaque and/or chimpanzee iPSCs were dissociated to single-cells, resuspended in iPSC culture medium supplemented with 2 nM of TZV, and keep on ice. After transportation (~2-3 hours in portable incubator), the E5 rhesus macaque blastocysts were transferred to warm M2 medium (CosmoBio, Carlsbad, USA) under a microscope. A piezo-driven micromanipulator (Prime Tech, Tokyo, Japan) was used to drill through the zona pellucida and trophectoderm, after which approximately 10 iPSCs were introduced into the blastocoel. The embryos were then washed 3 times in embryo media (see *Rhesus macaque oocyte retrieval, fertilization and embryo culture*), after which they were placed in a 70 μL drop containing a 1:1 ratio of embryo media and N2B27 basal media (*5*). Images where captured with Operetta (PerkinElmer, USA).

#### Differentiation of iPSC into cardiomycytes

iPSCs were differentiated into iPSC derived cardiomyocytes (iPSC-CMs) using a 2D monolayer differentiation protocol and were maintained in a 5% CO_2_/air environment(*35, 36*). Briefly, iPSC colonies were dissociated with 0.5 mM EDTA (Gibco) into single-cell suspension and re-suspended in E8 media containing 10 μM Rho-associated protein kinase inhibitor (Sigma). Approximately 100,000 cells were re-plated into Matrigel-coated 6-well plates. iPSCs were next cultured to 80-90% cell confluence, and then treated for 2 days with 6 μM CHIR99021 (Selleck Chemicals) in RPMI/B27 supplement without insulin to activate WNT signaling and induce mesodermal differentiation. On day 2, CHIR99021 was removed and cells were cultured in RPMI/B27 medium in the absence of insulin (*35, 37*). On day 3, cells were treated with 5 μM IWR-1 (Sigma) to inhibit WNT pathway signaling and further promote cardiogenesis. On day 7, IWR-1 was removed from the medium and cells were placed in RPMI/B27 in the absence of insulin. From day 7 onwards, cells were placed on RPMI+B27 supplemented with insulin until beating was observed. At this point, cells were treated in starvation medium (RPMI/B27 in the absence of insulin and glucose) for 3 days. Following purification via starvation medium, the survived cells were cultured in RPMI/B27 supplemented with insulin. Starvation was repeated at day 20 for a period of three days. When re-plating iPSC-CMs for downstream use, cells were dissociated with 10X TrypLE (Life Technologies) into a single-cell suspension and seeded on Matrigel-coated plates.

